# Dual UTR-A novel 5′ untranslated region design for synthetic biology applications

**DOI:** 10.1101/775643

**Authors:** Simone Balzer Le, Ingerid Onsager, Jon Andreas Lorentzen, Rahmi Lale

## Abstract

Bacterial 5′ untranslated regions of mRNA (UTR) involve in a complex regulation of gene expression; however, the exact sequence features contributing to gene regulation are not yet fully understood. In this study, we report the design of a novel 5′ UTR, dual UTR, utilising the transcriptional and translational characteristics of 5′ UTRs in a single expression cassette. The dual UTR consists of two 5′ UTRs, each separately leading to either increase in transcription or translation of the reporter, that are separated by a spacer region, enabling *de novo* translation initiation. We rationally create dual UTRs with a wide range of expression profiles and demonstrate the functionality of the novel design concept in *Escherichia coli* and in *Pseudomonas putida* using different promoter systems and coding sequences. Overall, we demonstrate the application potential of dual UTR design concept in various synthetic biology applications ranging from fine-tuning of gene expression to maximisation of protein production.

## Introduction

The DNA region corresponding to the 5′ untranslated region of mRNA (UTR) plays a central role in gene and protein expression. At the DNA level, it is involved in transcript formation due to the interplay between promoter and the initially transcribed sequences (ITS), which covers the first 15 nt in *Escherichia coli*^1–3^. At the mRNA level, it influences transcript stability and translation because of secondary structure formation^4–6^, interaction with external factors i.e. proteins^7^, metabolites^8, 9^ and short RNAs^10^ in addition to ribosome binding and translation initiation^11^. The translation initiation mainly involves around 15 nt preceding the start codon^12^ including the Shine-Dalgarno (SD) sequence as well as the 5′ end of the following coding sequence^4^. In bacteria, transcription and translation are coupled and a physical link between transcript formation and transcript turnover (translation and mRNA degradation) has been shown^13^. It has been reported that the translation rate is likely to affect the transcription rate which indirectly affects mRNA stability^14–17^. The DNA sequence of the 5′ UTR has also an under-recognised role on transcript formation involving nucleotides downstream of the ITS^18, 19^. With all the above mentioned characteristics, 5′ UTR is one crucial contributor to the maintenance of a fine balance between transcription, transcript stability, and translation.

In synthetic biology applications, it is desirable to have a predictive control over the levels of gene expression^20–22^. Currently, there are several *in silico* tools available enabling the design of synthetic 5′ UTR sequences for efficient translation initiation^15, 23–25^, and these are also applied in combination with a selection of promoters^15, 22, 26^. However, because of 5′ UTRs sequence proximity both to promoter and coding sequence regions a physical context dependency exists^27^, and it has consequently proven difficult to design optimal 5′ UTR sequences solely based on translational properties^15^. The multiple functionalities overlapping in 5′ UTRs, hence represents an optimisation challenge for modular design of biological circuitry.

We previously constructed a 5′ UTR plasmid DNA library in which a 22 nt stretch, of the 32 nt long 5′ UTR DNA sequence upstream of a β-lactamase coding sequence was randomly mutagenised using synthetic degenerate oligos^18^. *E. coli* cells harbouring the plasmid DNA library was screened for clones that showed increased ampicillin resistance at the induced state (β-lactamase production positively correlates with the host’s ampicillin resistance) with each clone carrying a unique 5′ UTR variant. *E. coli* clones could be categorised into two distinct groups based on their increased ampicillin resistance phenotype: one group of clones with increased reporter translation, high translation rates per reporter transcript; and another group of clones with increased reporter transcription phenotype but not corresponding translation, low translation rates per reporter transcript^18^. The 5′ UTR variants were identified as a result of screening efforts using a single 5′ UTR, and the screening would not allow the identification of specific 5′ UTRs with respect to the reporter transcription or translation rate. With these observations we speculated that the DNA sequence composition of a single 5′ UTR leads to a compromise between transcriptional and translational processes; hence it might not be possible to identify a sequence composition that is optimised for both of these processes in a short 5′ UTR sequences.

In this study, we designed new functional screening tools (bicistronic artificial operons) to identify 5′ UTR variants with desired transcriptional and translational characteristics. We constructed dual UTRs by incorporating the identified 5′ UTRs in a single expression cassette. We then demonstrate that the combination of 5′ UTRs with transcriptional and translational characteristic can have a synergistic effect on the gene and protein expression beyond the expression levels achieved with single 5′ UTRs. We demonstrate the functionality of the dual UTR concept in *Escherichia coli* and in an emerging SynBio chasis *Pseudomonas putida*^28^.

## Results

### Design, construction and screening of artificial operon plasmid DNA libraries

Initially, we sought to determine whether it would be possible to identify 5′ UTR variants that specifically lead either to increased reporter gene or protein expression. We designed two (bicistronic) artificial operons: pAO-Tr and pAO-Tn (p: plasmid; AO, artificial operon; Tr, transcription; Tn, translation), to identify 5′ UTR sequences that specifically lead to increased expression of β-lactamase at the mRNA and protein levels, respectively. Both artificial operons harbour the phosphoglucomutase encoding sequence, *celB*, as the first gene; a spacer region; and the β-lactamase encoding sequence, *bla*, as the second gene (Figure 1). The *celB* gene was chosen as it can be efficiently transcribed and translated, hence would not introduce any undesired restrictions^5^; and the *bla* gene was chosen as host’s resistance to ampicillin correlates with the produced amounts of β-lactamase, simplifying the identification of clones with the desired phenotype^18, 29^. As for the spacer region, it ensures that the translation of *bla* only occurs through *de novo* initiation as opposed to translational read-through^5^. The occurrence of *de novo* initiation was confirmed by eliminating the SD sequence upstream of *bla*, by replacing the SD sequence GGAG with CCTC, that led to abolished β-lactamase expression (results not shown). Expression in both artificial operons are driven by the positively regulated XylS/*Pm* regulator/promoter system, and a plasmid with the broad-host range mini-RK2 replicon is used as a vector in both constructs^30^.

**Figure 1.**
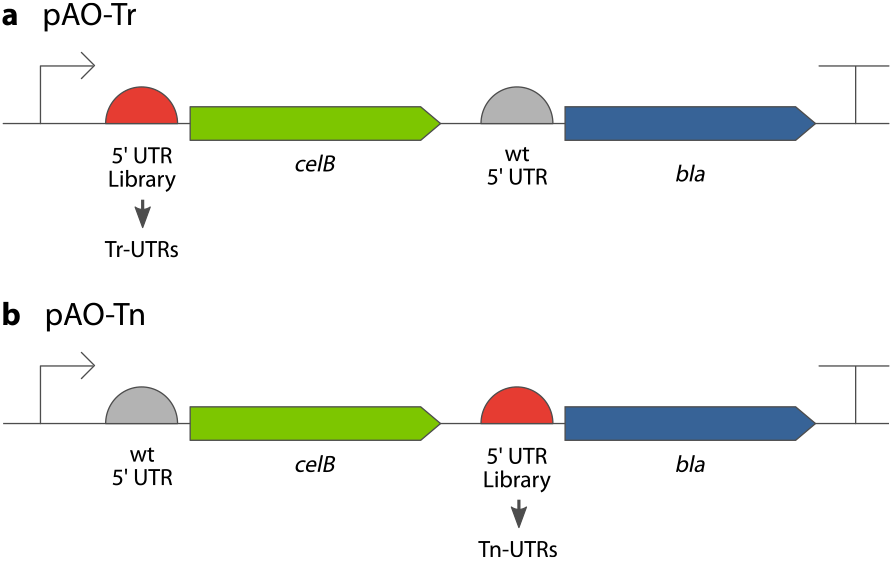
Composition of the artificial operons in plasmids pAO-Tr and pAO-Tn. The inducible XylS/Pm promoter system (indicated by an arrow) drives the expression in both operons, which consists of a 5′ UTR; a phosphoglucomutase encoding sequence, *celB*; a spacer region; a 5′ UTR; and a β-lactamase encoding sequence, *bla*. Two 5′ UTR plasmid DNA libraries were constructed using both artificial operons, and the 5′ UTR variants identified from pAO-Tr were named Tr-UTRs (a), and 5′ UTR variants identified in pAO-Tn were named Tn-UTRs (b). Both artificial operons have transcription terminators on their both ends (indicated by a T). p, plasmid; AO, artificial operon; Tr, transcription; Tn, translation.

Next, a 34 nt long oligo, with 22 nt degenerate nucleotide composition, was used for the construction of two 5′ UTR plasmid DNA libraries. For the transcriptional screening, using pAO-Tr (plasmid maps are provided as supplementary materials), a plasmid DNA library was constructed by cloning the degenerate oligo upstream of the first gene, *celB* (Figure 1a). Any observed increased expression of β-lactamase (detected as increased ampicillin resistance) would be as a consequence of increased transcription due to the presence of a particular 5′ UTR variant. For the translational screening, using pAO-Tn, the same degenerate 5′ UTR oligo was cloned upstream of the second gene, *bla* (Figure 1b). Any increased ampicillin resistance observed among the library clones would be as a consequence of increased *de novo* translation of the *bla* gene.

Finally for the screening, the recombinant *E. coli* clones harbouring the 5′ UTR plasmid DNA libraries (~280,000 and ~370,000 clones, in pAO-Tr and pAO-Tn, respectively) were plated on agar media containing 0.1 mM *m*-toluic acid (induces transcription from Pm) and ampicillin (ranging from 0.5 to 3.5 g/L). Multiple *E. coli* clones were isolated with increased ampicillin resistance phenotype, hypothesised to be either due to increased transcription (led to identification of Tr-UTR variants), or as a consequence of increased translation (led to identification of Tn-UTR variants). Identified clones could grow up to 2.5 g/L ampicillin in the presence of 0.1 mM *m*-toluic acid. To determine the 5′ UTR DNA sequences that led to increased ampicillin resistance several clones were selected, plasmid DNAs isolated and sequenced. Among the identified clones (Supporting Online Material [SOM] Table S1) three Tr-UTRs r31, r36, r50 (Figure 2a), and four Tn-UTRs n24, n44, n47, n58 (Figure 2b) were randomly selected for further characterisation along with the wildtype (wt) 5′ UTRs, and a previously identified 5′ UTR variant, LV-2^18^ (leading to increased transcription), summing up to five 5′ UTRs in each category. To ensure that the initially observed ampicillin resistance levels were solely caused by the mutations within the identified 5′ UTRs, synthetic oligonucleotides harbouring the identified mutations were synthesised and cloned back into pAO-Tr and pAO-Tn, and their ampicillin resistance phenotype were determined (Figure 2c). With the above mentioned screening and ampicillin resistance characterisations, we concluded that the artificial operons were suitable as a screening tool enabling the identification 5′ UTR variants that specifically lead to either increased gene or protein expression.

**Figure 2.**
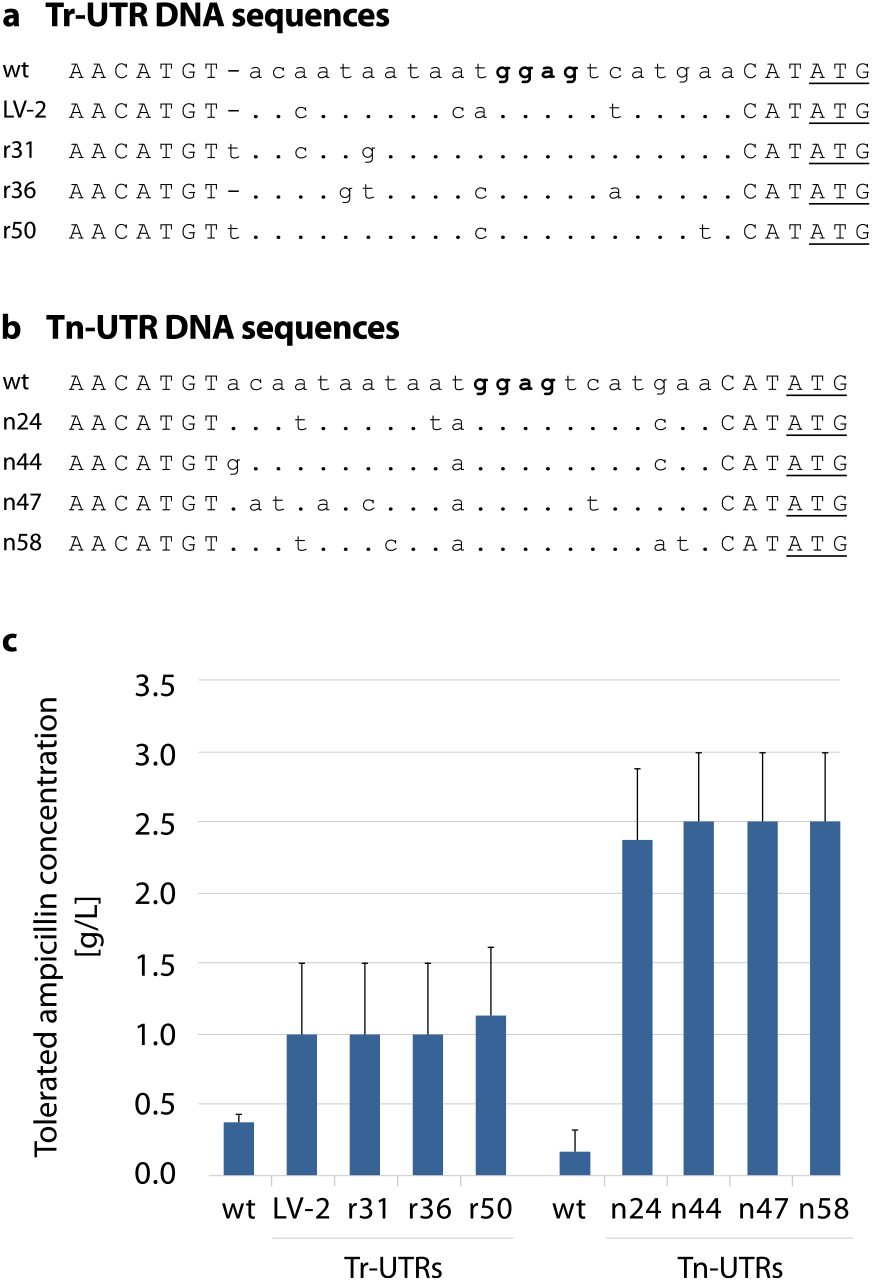
Nucleotide sequence composition and the phenotype of *E. coli* clones harbouring plasmids with different Tr- and Tn-UTRs. 5′ UTR DNA variants were identified by screening pAO-Tr- and pAO-Tn-based 5′ UTR libraries for increased ampicillin resistance. The 5′ UTR variants r31, r36, r50 are identified from the Tr-UTR library (a); while n24, n44, n47, n58 from the Tn-UTR library (b). The 5′ UTR variant LV-2 serves as an internal control that was previously shown to be leading to increase in transcription of the reporter^18^. Identical nucleotides are indicated by dots and point mutations are indicated with letters. Nucleotides that are not mutagenised are typed in capital letters including the PciI (ACATGT) and NdeI (CATATG) recognition sequences. The putative SD sequence (ggag) is highlighted in boldface. The ATG start codon (part of the NdeI site) is underlined. The ampicillin resistance phenotypes of the *E. coli* clones harbouring Tr- or Tn-UTR DNA sequences were characterised at the induced state with 0.1 mM m-toluic acid (c). This low concentration was used to make sure that resistance levels were in a range allowing us to distinguish moderate phenotypic differences among the clones. Results are presented as averages of the highest ampicillin concentrations at which growth was observed. Error bars point to the next tested ampicillin concentration at which no growth was observed.

### Design and construction of dual UTR

The screening of the artificial operons gave us access to unique 5′ UTR variants. Now having these 5′ UTRs, we took a rational design approach and systematically combined them with the objective of benefiting from both the transcriptional and translational characteristic of 5′ UTRs in a single dual UTR. A dual UTR consists of two unique 5′ UTRs separated by a spacer region: the first 5′ UTR proximal to the promoter is where we placed 5′ UTRs originating from the transcriptional screening (Tr-UTR); the second 5′ UTR is where we placed the 5′ UTRs identified from the translational screening (Tn-UTR); while the spacer region provides enough space for physical separation of mutations affecting transcription and translation (Figure 3a, SOM Figure S1). With this design concept, we constructed 25 dual UTRs (Figure 3b). In all 25 constructs, the dual UTRs were cloned downstream of the *Pm* promoter, and the entire expression cassette was resting on a plasmid with the broad-host range mini-RK2 replicon.

**Figure 3.**
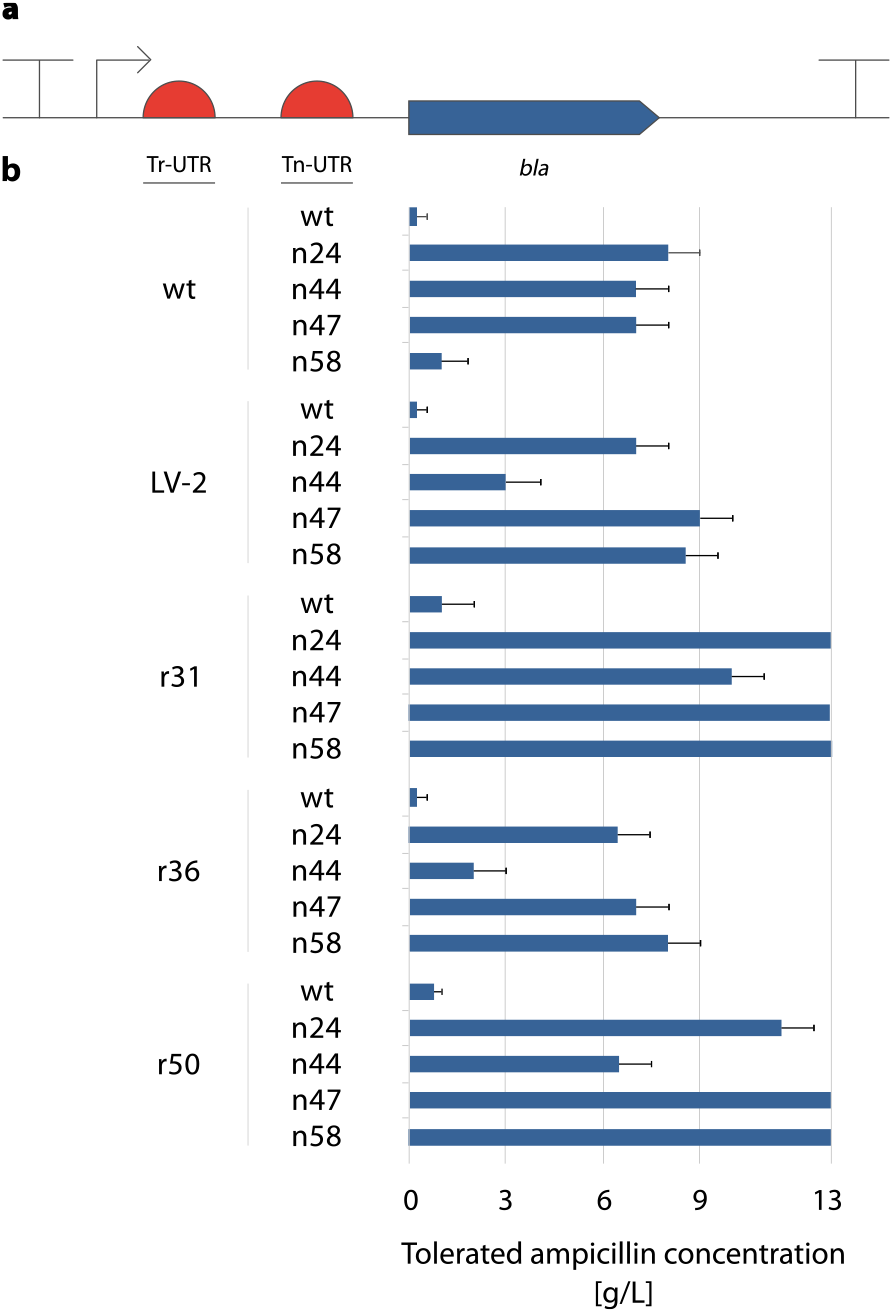
Schematic view and characterisation of ampicillin resistance phenotype in *E. coli* clones harbouring plasmids with different dual UTR constructs. The inducible XylS/*Pm* promoter system (indicated by an arrow) drives the expression in all constructs. The dual UTR consists of a Tr-UTR, a spacer region and a Tn-UTR (a). The ampicillin resistance phenotype of the *E. coli* clones harbouring each of the 25 dual UTR combinations were characterised using 2 mM *m*-toluic acid to provide full induction (b). Thirteen g/L ampicillin was the highest concentration tested. Error bars point to the next tested ampicillin concentration at which no growth was observed.

### Quantification of effects on gene and protein expression

Initially, the ampicillin resistance phenotype of the *E. coli* clones each harbouring one of the 25 dual UTRs were characterised (Figure 3b). The observed ampicillin resistance levels indicated that all the dual UTR constructs were leading to increased reporter gene and protein expression as compared to expression levels reached with the wtwt dual UTR construct. A striking finding was that the observed ampicillin resistance phenotypes of the *E. coli* clones containing the dual UTR constructs were much higher than the resistance levels observed with the *E. coli* clones containing the individual Tr- and Tn-UTRs on their own (Figure 2c). This initial ampicillin resistance characterisations demonstrated that with the dual UTR design concept strong gene expression and high protein production can be achieved. It was also encouraging to observe that high expression levels could be reached by the combination of five Tr- and Tn-UTRs only.

Next, we sought to determine the reporter transcript and protein levels resulting from the dual UTR constructs. Four *E. coli* clones each harbouring one of the dual UTR constructs were grown under induced state and the transcript levels, by relative quantitative real-time reverse-transcription PCR (qPCR); and the enzymatic activities, by β-lactamase enzymatic activity assays, were determined (Figure 4a). The results obtained were in agreement with the design concept that r31, in r31wt dual UTR construct, led to increase in reporter transcript (despite moderate); while n47, in wtn47, led to 10-fold increase in *bla* transcript and a 42-fold increase in β-lactamase enzyme activity compared to the levels of expression reached with the wtwt dual UTR construct. Strikingly, the r31n47 dual UTR led to a high reporter gene and protein expression, 46- and 170-fold increase respectively, compared to the expression levels observed with the wtwt dual UTR construct. For quantitative comparison, we also calculated the translation efficiency for each construct by simply determining the protein-to-transcript ratio. Translation efficiencies were 1, 0.6, 4.2, and 3.7 for the wtwt; r31wt; wtn47; and r31n47 dual UTR constructs, respectively. The observed translation efficiencies were in agreement with the dual UTR design concept that while r31 was leading to increased reporter transcription, hence lower translation efficiency; the n47 was leading to increased reporter translation, hence higher translation efficiency. Taken together, the increased expression observed with the r31n47 dual UTR construct was the result of synergistic effect of 5′ UTRs with transcriptional and translational characteristics.

**Figure 4.**
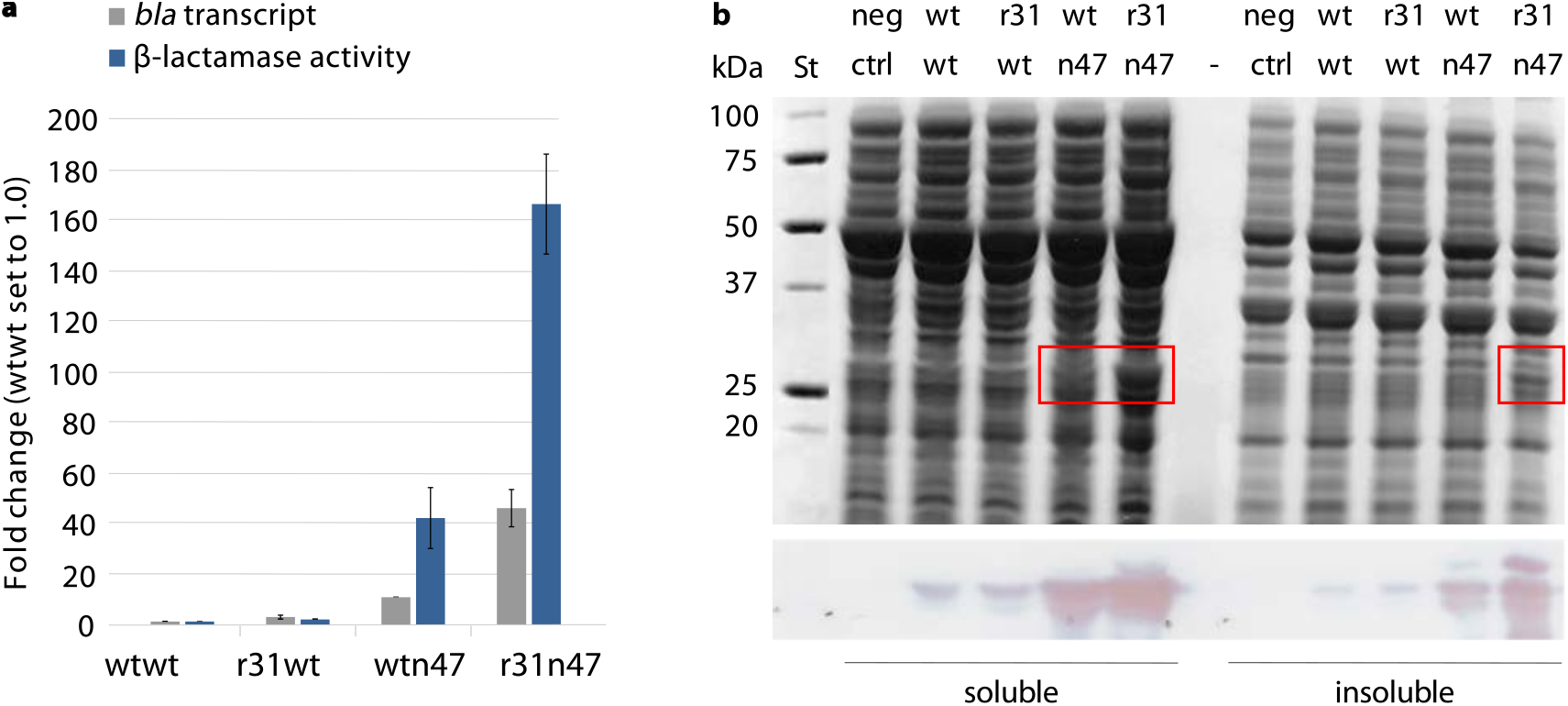
Quantification of *bla* expression levels in *E. coli* clones harbouring plasmids with different dual UTR constructs. (a) Relative *bla* transcript amounts and β-lactamase enzymatic activities (values for the wtwt dual UTR construct were arbitrarily set to 1.0) were determined after 5 hours of growth at the induced state with 2 mM m-toluic acid. (b) SDS-PAGE (top) and Western blot (bottom) images of the soluble and insoluble protein fractions of *E. coli* clones producing β-lactamase. Visible β-lactamase bands are highlighted with red rectangles. Error bars indicate the standard deviations calculated from three replicates. St: Precision Plus Dual Color Protein standard (Bio-Rad); neg ctrl: plasmid-free strain.

Next, the total cellular protein production was assessed by SDS-PAGE. β-lactamase could be visualised on the gel in the soluble fraction of sonicated cell lysates from clones harbouring constructs with the wtn47 and r31n47 dual UTRs. The enzyme could also be visualised in the insoluble fraction from the r31n47 dual UTR construct (Figure 4b, upper panel). Specific detection of β-lactamase was also performed by Western blotting and the signal strengths correlated with the quantified enzymatic activities (Figure 4b, lower panel).

### Phenotypic quantification with a red fluorescent protein

The Tn-UTR variants used in the dual UTR constructs were identified using the *bla* gene; therefore we were interested in testing to what extent the observed increased reporter gene expression was coding sequence (CDS) specific. To assess the potential context dependency, the β-lactamase CDS was substituted with mCherry CDS, encoding for a red fluorescent protein, in wtwt, r31wt, wtn47 and r31n47 dual UTR constructs and mCherry transcript levels, fluorescent intensities and total proteins were quantified. Here we also calculated the translation efficiencies resulting from the four dual UTR constructs and the ratios were: 1, 0.5, 1.2, and 1.4 for the wtwt, r31wt, wtn47 and r31n47 dual UTR constructs, respectively. The phenotypic quantification with mCherry also indicate that r31 leads to increase in reporter transcription, whereas the n47 leads increase in reporter translation (Figure 5a). Correspondingly, the mCherry protein could also be easily visualised on an SDS-PAGE, both in the soluble and insoluble fraction, particularly from the r31n47 dual UTR construct (Figure 5b). These results indicate that the observed increase in expression was not CDS-specific. This finding indicates the suitability of the novel 5′ UTR design concept for applications aiming for high levels of expression for instance heterologous protein production in bacteria.

**Figure 5.**
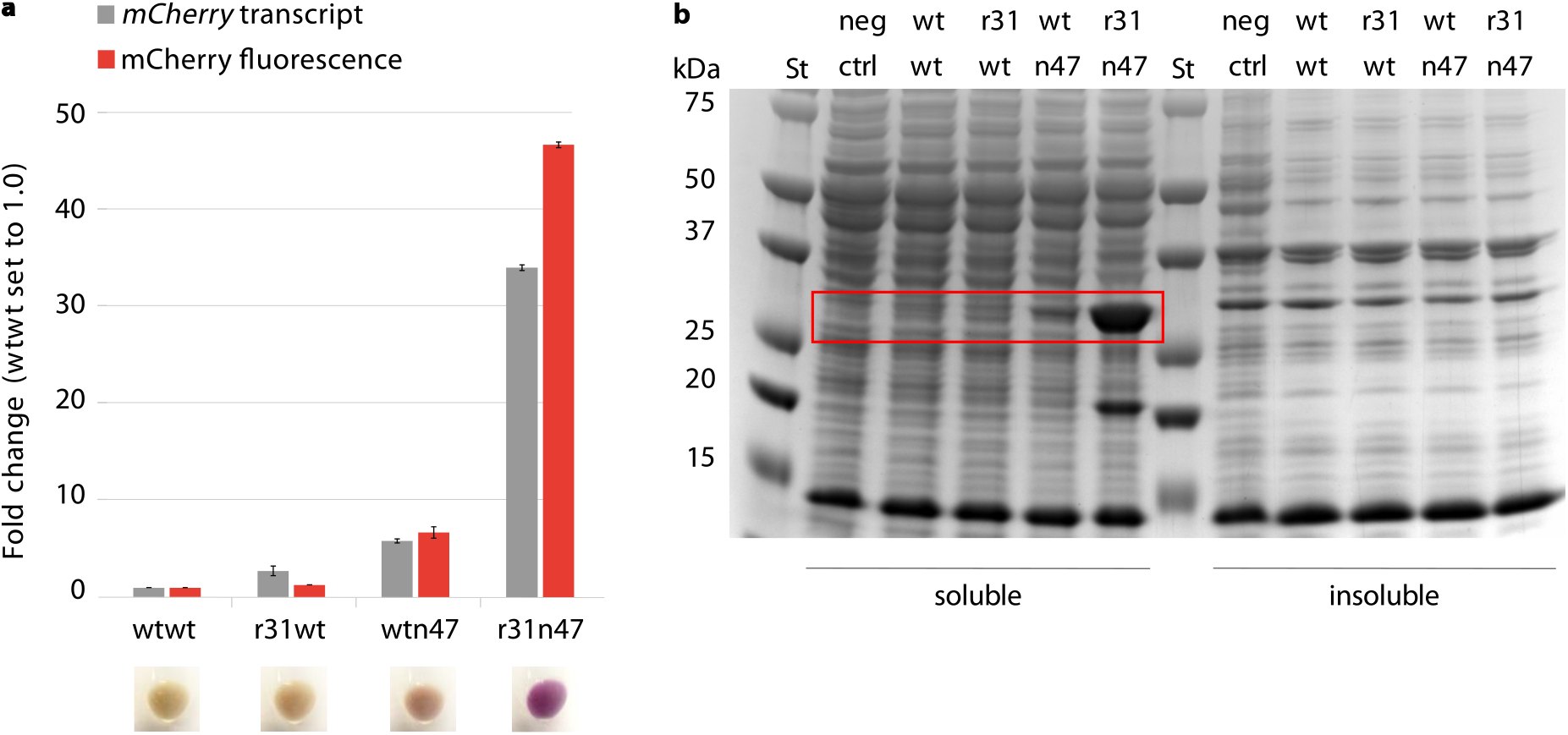
Quantification of *mCherry* expression levels in *E. coli* clones harbouring plasmids with different dual UTR constructs. (a) Relative *mCherry* transcript amounts and mCherry fluorescent intensities were quantified after 5 hours of growth at the induced state with 2 mM *m*-toluic acid. Fluorescence intensities were determined directly from the cultures and normalised with OD_600_ values. Values for the wtwt dual UTR construct were arbitrarily set to 1.0. The four images below the panel a shows bacterial pellets obtained from the four cultures indicated. Error bars indicate the standard deviations calculated from three replicates. (b) SDS-PAGE image of *E. coli* clones producing mCherry. Visible mCherry protein bands are highlighted with a red rectangle. St: Precision Plus Dual Color Protein standard (Bio-Rad); neg ctrl: plasmid-free strain.

### *In silico* design of Tn-UTRs

The results reported so far indicate that a combination of 5′ UTRs in a dual UTR can lead to strong gene and protein expression, at least for the two tested selection marker and reporter protein, *bla* and mCherry in *E. coli*, respectively. However, context dependency between 5′ UTR sequences and CDS cannot be excluded if more genes are to be tested^27^. Therefore we applied a widely used *in silico* tool, the RBS calculator^24^, in designing Tn-UTRs that are specifically designed for the gene of interest to achieve high translation initiation rates (TIR, SOM Table S2). In the design, the *bla* and *mCherry* genes were used as reporters, and *E. coli* was chosen as the host. In total, six such designed Tn-UTRs were synthesised: three for the β-lactamase and three for the mCherry CDSs (named as dTn-UTRs). The predicted TIR values were 60-to 80-fold higher compared to n47-bla, and 50-to 70-fold higher compared to n47-*mCherry* (SOM Table S3). All six dTn-UTRs were cloned into the dual UTR either with the wt or the r31 at the Tr-position (Figure 6). The phenotypic characterisations revealed that all three dTn-UTRs with the *bla* gene led to a similar increase in expression levels achieved with the n47 combination (Figure 6a). Similar observations were also made for the construct with the *mCherry* gene: Tr-UTR r31 in combination with the dTn-UTRs led to 9- (dTn4) and 46-fold (dTn5 and dTn6) relative increase vs. 58-fold with the Tn-UTR n47 (Figure 6b). These findings indicate that designed 5′ UTRs can also be used in the dual UTR context, and even higher expression levels, beyond the predicted levels, can be achieved by the combination of designed Tn-UTRs with Tr-UTRs.

**Figure 6.**
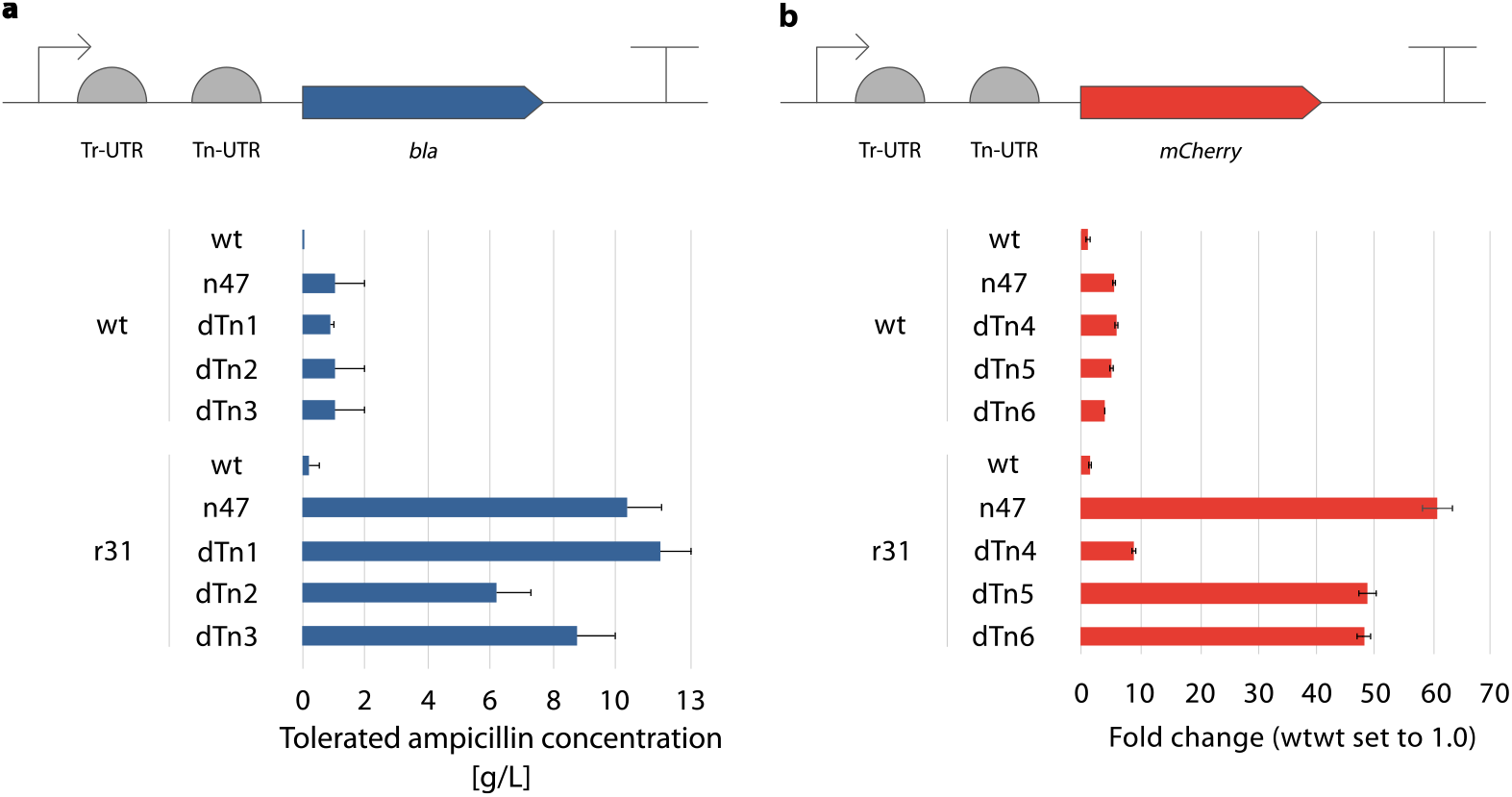
Phenotypic characterisation of the dual UTR constructs with *in silico* designed Tn-UTRs. (a) The ampicillin resistance phenotype of the *E. coli* clones harbouring each of the ten dual UTR combinations were characterised at the induced state with 0.1 mM m-toluic acid. The inducible XylS/Pm promoter system (indicated by an arrow) drives the expression in both constructs. Error bars point to the next tested ampicillin concentration at which no growth was observed. (b) mCherry fluorescent intensities were quantified after 5 hours of growth at the induced state with 2 mM *m*-toluic acid. Fluorescence intensities were determined directly from the cultures and normalised with OD_600_ values. The fluorescent intensity value for the wtwt dual UTR construct was arbitrarily set to 1.0. Error bars indicate the standard deviations calculated from three replicates.

### Functionality assessment in an alternative host

The broad-host range mini-RK2 replicon and the expression system XylS/*Pm* are both known to function in many Gram-negative bacteria^31^ including *Psuedomonas* species. The emerging SynBio chasis, *P. putida*^28^, has the same anti-SD sequence within its 16S rRNA as *E. coli*. We therefore sought to assess the functionality of the dual UTR concept in *P. putida* strain KT2440. The dual UTR constructs wtwt, r31wt, wtn47and r31n47 with the *mCherry* CDS were transferred to *P. putida* and the mCherry fluorescent intensities were quantified (Figure 7). The strong synergistic effect seen with dual UTR constructs in *E. coli* was also observed in *P. putida*. The relative expression levels reached among the dual UTR constructs were weaker in *P. putida* than the observed relative expression levels in *E. coli*. The reason for the relative weakness appeared to be due to the reference construct, wtwt, leading to expression of mCherry at much higher levels in *P. putida* than in *E. coli* as judged by the stronger bands visualised on the SDS-PAGE gel (Figure 7b). The results obtained in *P. putida* indicates that the functionality of the dual UTR concept is not only limited to *E. coli*; hence in principle it can be used in a wide range of bacterial hosts.

**Figure 7.**
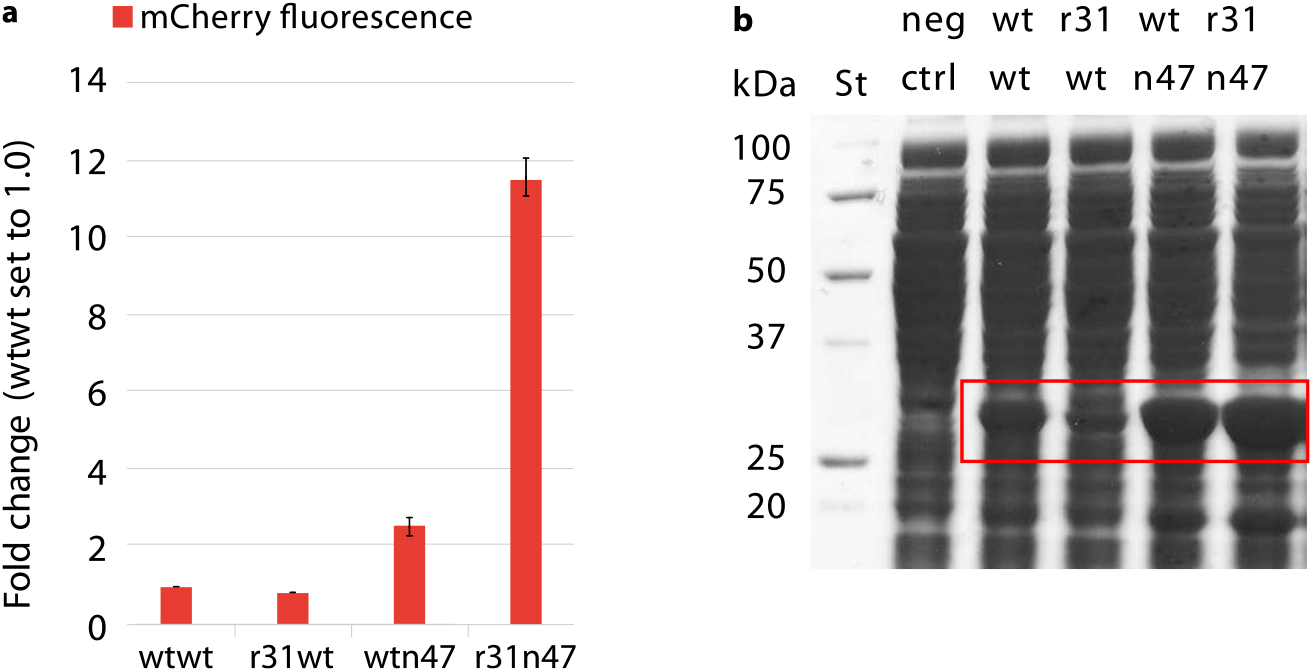
Quantification of *mCherry* expression levels in *P. putida* clones harbouring plasmids with four different dual UTR constructs. (a) Relative mCherry fluorescent intensities were quantified after 5 hours of growth at the induced state with 2 mM *m*-toluic acid. Fluorescence intensities were determined directly from the cultures and normalised with OD_600_ values. The fluorescent intensity value for the wtwt dual UTR was arbitrarily set to 1.0. Error bars indicate the standard deviations calculated from three replicates. (b) SDS-PAGE image of *P. putida* clones producing mCherry. Visible mCherry protein bands are highlighted with a red rectangle. St: Precision Plus Dual Color Protein standard (Bio-Rad); neg ctrl: plasmid-free strain.

### Assessment of the positional effect

The conceptualisation of dual UTR was the result of a rational design approach. The particular 5′ UTRs were rationally placed in dual UTR given their transcriptional and translational characteristics which led to the synergistic effect resulting in high reporter gene and protein expression. Observing the synergistic effect on gene expression, we sought to determine whether the strong expression levels could also be achieved irrespective of the position of the 5′ UTRs in dual UTR. To assess the positional effect on the observed phenotype, we constructed dual UTRs by placing the Tn-UTRs n47 and n58 in Tr-position, and the Tr-UTRs r31 and r50 in Tn-position. After the construction of the plasmids with TnTr dual UTR combinations we quantified the resulting fluorescent intensities under induced and uninduced states in *E. coli* (Figure8). It was intriguing to observe that the synergistic effect observed with dual UTRs were position dependent as the dual UTRs with TnTr combinations were not leading to any significant expression at all.

**Figure 8.**
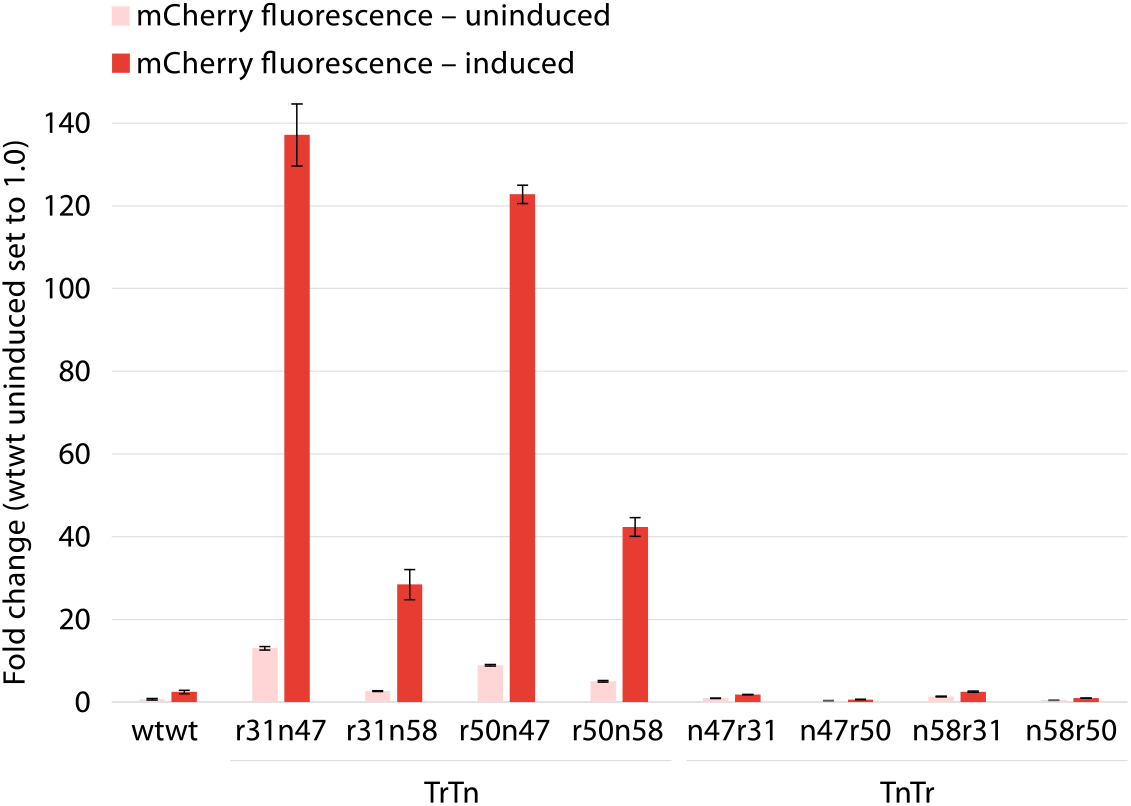
Quantification of *mCherry* expression levels in *E. coli* clones harbouring plasmids with TrTn and TnTr dual UTR constructs. Relative mCherry fluorescent intensities were quantified after 5 hours of growth at the induced state with 2 mM m-toluic acid. Fluorescence intensities were determined directly from the cultures and normalised with OD_600_ values. The fluorescent intensity value for the wtwt dual UTR construct at the uninduced state was arbitrarily set to 1.0. Error bars indicate the standard deviations calculated from three replicates.

**Figure 9.**
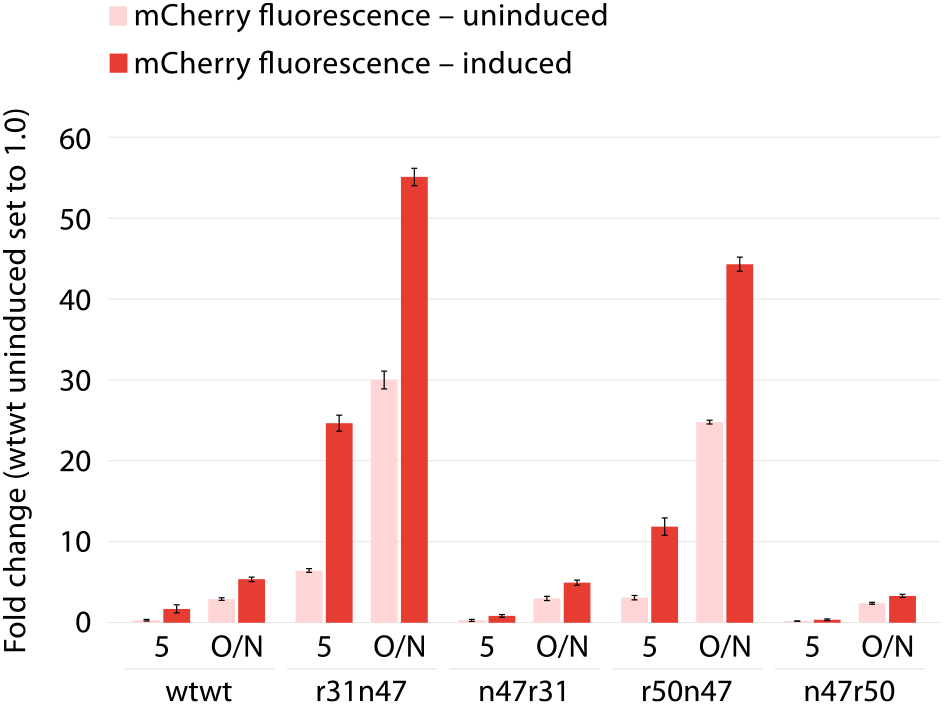
Quantification of *mCherry* expression levels under the control of the AraC/*P_BAD_* promoter system in *E. coli* clones harbouring plasmids with TrTn and TnTr dual UTR constructs. Relative mCherry fluorescent intensities were quantified after 5 hours and 18 hours of overnight (O/N) growth at the induced state with L-arabinose to a final concentration of 0.2%. Fluorescence intensities were determined directly from the cultures and normalised with OD_600_ values. The fluorescent intensity value for the wtwt dual UTR construct at the uninduced state was arbitrarily set to 1.0. Error bars indicate the standard deviations calculated from three replicates.

### Functionality assessment with an alternative promoter system

Observing that the dual UTR was functional with another coding sequence, in another host, and the correct positioning of the 5′ UTRs was necessary for the observed synergistic effect, we questioned whether the observed increase in gene and protein expression with dual UTRs was due to an emergent property of the *Pm* promoter. For this functionality assessment, we chose to substitute the XylS/*Pm* with the AraC/*P_BAD_* system in dual UTR constructs, both for the original design of TrTn as well as TnTr combinations. We characterised the phenotype of the *E. coli* clones by quantifying the fluorescent intensities resulting from each construct under induced and uninduced states after 5 and 18 hours (O/N) of growth. The results of this experiment made it clear that the synergistic effect observed in dual UTRs was not an emergent property of the *Pm* promoter as similar expression levels were also reached when we used the *P_BAD_* promoter in combination with dual UTRs, both for the TrTn and TnTr combinations. The dual UTR constructs, however, led to much higher background expression from the *P_BAD_* promoter in the absence of induction as compared to the expression levels achieved with the *Pm* promoter.

### Transcript stability and translational arrest

It has been reported that high translation rate (high ribosome occupancy) occurring on mRNA leads to protection of the mRNA against nucleotide degradation^14–17^. Given the high reporter transcript levels observed with the dual UTR constructs, we sought to determine the decay rates of the reporter transcripts originating from the dual UTR constructs, and relative presence of reporter transcripts under translational arrest. For determining the reporter transcript stability, we performed an inducer wash-out assay^32^. The wash-out assay relies on the passive diffusion characteristic of the *m*-toluic acid, inducer of the XylS/*Pm* system. *E. coli* clones were grown for 5 hours under induced state with 2 mM *m*-toluic acid and after this period of growth the growth medium was replaced with a fresh medium not containing the inducer, and five samples were taken with 2 minutes intervals. qPCR was performed on the samples and the mRNA decay rates were calculated for the four different dual UTR constructs (Figure 10). The stability of the reporter transcript resulting from the dual UTRs appear to be correlated with the levels of protein expression for the wtn47 and r31n47 dual UTR constructs. While the observed lower stability for the reporter transcript from the r31wt does not correlate with the measured mCherry fluorescence intensities.

**Figure 10.**
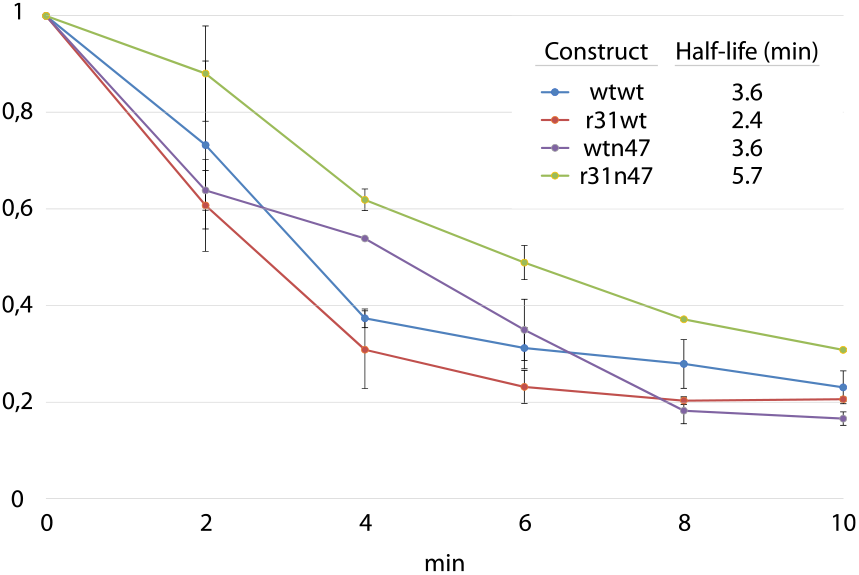
Quantification of mRNA decay rates in four dual UTR constructs in *E. coli. E. coli* clones were first grown for 5 hours under induced state with 2 mM *m*-toluic acid, and afterwards the growth medium was replaced with a fresh medium not containing the inducer. In total five samples were taken with 2 minutes intervals ranging from 0 to 10 minutes. qPCR analyses was performed on the samples and the mRNA decay rates were calculated for the four different dual UTR constructs.

Next, we performed an translational arrest experiment by the addition of chloramphenicol to the growth medium that leads to peptidyl transferase inhibition of bacterial ribosome, hence hinders amino acid chain elongation. For this experiment the transcript and fluorescence intensities were measured under two conditions: (i) by the induction of expression only with 2 mM *m*-toluic acid, to serve as a control (I); (ii) by the addition of chloramphenicol to a final concentration of 100 μg/mL and followed by induction of expression with 2 mM m-toluic acid, to assess the accumulation of reporter transcript in the absence of translation (CI) (Figure 11a). The reporter transcript accumulation resulting from the r31n47 dual UTR construct is not negatively affected by the blocking of translation as the transcript levels remain similar as compared to the condition I (Figure 11b). Hence the observed transcript stability, determined by the decay rate experiment, is not due to the high translation observed in the r31n47 dual UTR construct. This same observation also holds true for the wtn47 dual UTR construct. One puzzling observation was the increased fluoresence accumulation resulting from the r31wt construct. The decay rate indicates that the reporter transcript is the least stable compared to the other three reporter transcripts resulting from the dual UTR constructs; however, r31wt still leads to protein accumulation in the presence of chloramphenicol. This observation points to the presence of an unknown biological process that still leads to protein accumulation in the presence of chloramphenicol.

**Figure 11.**
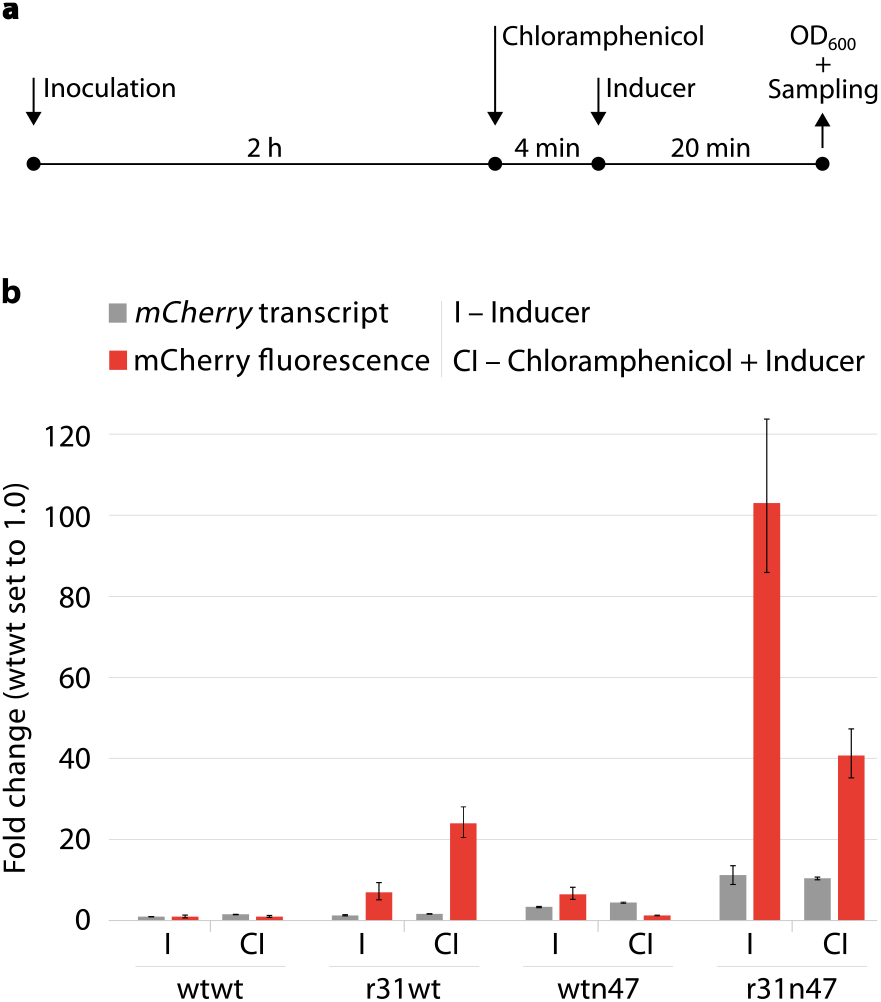
Quantification of the reporter transcript and mCherry fluorescence intensities under translational arrest. (a) The schematic view of the experimental set-up. *E. coli* clones with different dual UTR constructs were inoculated to a fresh medium and were grown for 2 hours. Afterwards it was either followed by induction of expression only with 2 mM *m*-toluic acid (I) or by the addition of chloramphenicol to a final concentration of 100 μg/mL and followed by induction of expression with 2 mM *m*-toluic acid (CI). The quantified relative *mCherry* transcript amounts and mCherry fluorescent intensities were depicted in panel b. Fluorescence intensities were determined directly from the cultures and normalised with OD_600_ values. Values for the wtwt dual UTR construct at the condition I were arbitrarily set to 1.0. Error bars indicate the standard deviations calculated from three replicates.

## Discussion

In this study, we present a novel 5′ UTR design named dual UTR. The dual UTR concept both relies and benefits from the role of 5′ UTRs in transcriptional and translational regulation in bacteria. For this study, we designed artificial operons to identify 5′ UTRs that individually lead to increase in reporter transcription or translation. The combination of 5′ UTRs in a single UTR led to the observation of a synergistic effect resulting in increased expression both at the level of reporter transcript and final protein. The strong gene expression levels achieved with dual UTRs point to the importance of the role of 5′ UTRs in transcriptional regulation. Based on the mRNA decay and translational arrest experiments we speculate that the reporter transcript accumulations in dual UTR are not dependent on the translation events. This points to an important finding that reporter transcripts can be generated, by an unknown mechanism, that is not as a consequence of protection due to high ribosome occupancy. Further research directions will be on the study of single gene expression events by labelled RNA polymerase, and ribosome profiling to understand the role and influence of 5′ UTRs in transcript accumulation, especially in relation to ribosome occupancy.

In synthetic biology applications 5′ UTRs are utilised due to their involvement in translational regulation only. Therefore the functional role of 5′ UTRs in transcriptional regulation has not been realised in the modular design of biological circuitry. In the biological circuitry design different levels of gene expression is achieved either by the use of native promoters, or variants thereof with different strength; or by the use of limited number of regulated promoters via differential induction levels. The use of plasmids with different copy-number has also been used as an another way of obtaining varied gene expression. Based on the experimental evidences reported in this study, we envision that the dual UTR concept can also be used as a way to achieve transcriptional regulation in biological circuitry design. This is especially relevant for studies using (unconventional) host microorganisms for which there is a limited number of characterised promoters exist.

The dual UTR design concept is also suitable for applications where the introduction of expression cassettes from plasmid-based systems (multiple copies) to chromosomes (single copy) suffers from the reduction in gene dosage. Hence dual UTRs can be used, even in combination with strong promoters, to achieve maximum transcription rates from a single expression cassette.

To conclude, a 5′ UTR involves in complex regulations both at the level of DNA and mRNA. The combination of Tr- and Tn-UTRs in dual UTR leads to a synergistic effect in gene and protein expression. The functionality of the dual UTR depends on the individual characteristics of the 5′ UTRs, and the correct positioning of Tr- and Tn-UTRs is crucial for the observed synergistic effect. The dual UTR as a concept is functional with different promoters and coding sequences both in *E. coli* and *P. putida*. With these demonstrations we anticipate that the concept described here are universally applicable to achieve a precise control of expression in bacteria for various synthetic biology applications.

## Methods

### Bacterial strains and growth conditions

Recombinant *E. coli* DH5*α* (Bethesda Research Laboratories), *E. coli* RV308 (ATCC 31608) and *P. putida* KT2440 were cultivated in Lysogeny Broth (LB) (10 g/L tryptone, 5 g/L yeast extract and 5 g/L NaCl) or on Lysogeny Agar (LB broth with 15 g/L agar) supplemented with 50 μg/mL kanamycin. Selection of *E. coli* DH5*α* transformants was performed at 37 °C, while 30 °C was used for all growth experiments. For the induction of the promoter systems, first overnight cultures were inoculated into fresh medium and were grown until the mid-log phase was reached, and afterwards were induced at the levels specified.

### DNA manipulations

Standard recombinant DNA procedures were performed as described by Sambrook and Russell^33^. DNA fragments were extracted from agarose gels using the QIAquick gel extraction kit and from liquids using the QIAquick PCR purification kit (QIAGEN). Plasmid DNA was isolated using the Wizard Plus SV Minipreps DNA purification kit (Promega) or the NucleoBond Xtra Midi kit (Macherey-Nagel). Synthetic oligonucleotides were ordered from Sigma-Aldrich or Eurofins. Restriction cloning was performed according to recommendations from New England Biolabs. PCR reactions were carried out with the Expand High Fidelity PCR System (Roche Applied Science) or Q5 High-Fidelity DNA Polymerase (NEB). *E. coli* clones were transformed using a modified RbCl protocol (Promega) and *P. putida* KT2440 was transformed using an electroporation protocol described by Hanahan and colleagues^34^. The constructed plasmids were confirmed by sequencing performed at Eurofins/GATC Biotech using primer 5′-AAC GGCCTGCTCCATGACAA-3′ for pAO-Tr-, pIB11-^18^, pdualUTR-; and primers 5′-CTTTCACCAGCGTTTCTGGGTG-3′ and 5′-CAAGGATCTTACCGCTGTTG-3′ for pAO-Tn-based constructs.

### Plasmid constructions

All plasmids are based on the broad-host range mini-RK2 replicon. For the construction of the pAO-Tr plasmid, the *bla* coding sequence was amplified from the plasmid pIB11^18^ with the primers 5′-GCAGGCGGAATTCTAATGAGGTCATGAACTTATGAGTATTCAACATT-3′ and 5′-CTAGAGGATCCCCGGGTACCTTTTCTACGG-3′, introducing the restriction sites EcoRI and BamHI, and was cloned into the pIB22^35^ plasmid as EcoRI-BamHI fragment downstream of the *celB* gene. This construction resulted in the plasmid pAO-Tr (SOM pAO-Tr.gb).

For the construction of pAO-Tn, the *celB* coding sequence and the DNA sequence corresponding to its 5′UTR were PCR amplified from the pAO-Tr plasmid using the primer pair 5′-ACCCCTTAGGCTTTATGCAACAgaaACAATAATAATGGAGTCATGAACtTATG-3′ and 5′-CTTTCACCAGCGTTTCTGGGTG-3′. The resulting PCR product was digested with Bsu36I and EcoRI and re-introduced into pAO-Tr using the same restriction sites leading to pAO-Tn(−1). By this cloning, the additional NdeI and PciI restriction sites were removed (indicated by small letters). The *bla* coding sequence was PCR amplified from pIB11 using the primer pair 5′-cggaattCAACATGTA CAATAATaatg-3′ and 5′-AGCTAGAGGATCCCCGGGTA-3′ and the resulting PCR product was cloned as an EcoRI-BamHI fragment into the pAO-Tn(−1) plasmid resulting in pAO-Tn (SOM pAO-Tn.gb).

For the individual characterisation of Tr- and Tn-UTR’s phenotype, 5′ UTR DNA sequences were integrated between the unique PciI and NdeI sites in the plasmid pIB11 as annealed pairs of forward and reverse synthetic oligonucleotides.

For the construction of plasmids carrying the dual UTRs, pdualUTR (SOM pdualUTR.gb), the native 5′ UTR DNA sequence upstream of the *bla* coding sequence in pIB11 was substituted with the annealed oligonucleotides 5′-CATGTACAATAATAATGGAGTCATGAACATATCTTCATGAGCTCCATTATTATTGTATATGTACAATAATAATGGAGTCATGAACA-3′ and 5′-TATGTTCATGACTCCATTATTATTGTACATATACAATAATAATGGAGCTCATGAAGATATGTTCATGACTCCATTATTATTGTA-3’.

For the construction of pdualUTR with the *mCherry* reporter gene an *E. coli* codon-optimized variant of the *mCherry* coding sequence (a gift from Yanina R. Sevastsyanovich, University of Birmingham) was PCR-amplified using primer pair 5′-GCTGCATATGGTTTCTAAAGGTGAAGAAG-3′ and 5′-GCTCGGATCCTTATCATTTATACAGTTCGTCCATAC-3′ and digested with NdeI and BamHI. The digested fragment was then used to replace the *bla* coding sequence in pdualUTR resulting in pdualUTR-mCherry (SOM pdualUTR-mCherry.gb).

For the construction of all dualUTR combinations annealed synthetic oligonucleotides flanked by PciI and SacI (Tr-dual UTR) and SacI and NdeI (Tn-dual UTR) sticky ends were inserted into pdualUTR or pdualUTR-mCherry using the respective restriction enzyme recognition sites.

For the construction of the *P_BAD_* system, the *P_BAD_* expression cassette was PCR amplified from pSB-B1b^36^ using primer pairs 5′-ATGGAGAAACAGTAGAGAGTTG-3’and 5′-TACATGGCTCTGCTGTAG-3’. Each of the five pdualUTR-mCherry vectors were PCR amplified using the reverse primer 5′-CTCCCGTATCGTAGTT ATC-3′ in combination with one of the forward primers 5′-CAACTCTCTACTGTTTCTCCATAACATGTTACCATGATAATGGAG-3′, 5′-CAACTCTCTACTGTTTCTCCATAACATGTTACAATAATAACGGAGTCATG-3′, 5′-CAACTCTCTACTGTTTCTCCATAACATGTAATAAACTAAAGGAGTTATG-3′ 5′-CAACTCTCTACTGTTTCTCCATAACATGTACAATAATAATGGAGTCATGAACATATC-3′ each targeting the dual UTR region in r31n47, r50n47, n47r31/n47r50, and wtwt respectively. The DpnI treated and purified PCR-products with overlapping ends were assembled using *in vivo* homologous recombination method in *E. coli* strain DH5*α*, leading to pdualUTR-PBAD-mCherry (SOM pdualUTR-PBAD-mCherry.gb). The correct constructs were confirmed by sequencing using the reverse primer binding to the mCherry coding sequence, 5′-GATGTCAGCCGGGTGTTTAAC-3′.

### Generation and screening of 5′ UTR libraries based on pAO-Tr and pAO-Tn

5′ UTR libraries were constructed in pAO-Tr and pAO-Tn by cloning the synthetic degenerate oligonucleotides between their respective NdeI and PciI restriction sites, as described previously^18, 30^. After transformation of plasmid DNAs to *E. coli* DH5*α*, libraries with ~280,000 transformants (pAO-Tr-based) and ~370,000 transformants (pAO-Tn-based) were generated.

### β-lactamase expression analysis

β-lactamase enzymatic assay was performed following the protocol described elsewhere^37^. For protein production analysis by SDS-PAGE or Western blot, *E. coli* RV308 was used. *E. coli* RV308 clones were grown in super broth (32 g/L peptone, 20 g/L yeast extract and 5 g/L NaCl). Expression was induced in the mid-log phase and cultures were harvested 5 hours after induction with 2 mM m-toluic acid. 0.1 g pellet (wet weight) were washed with 0.9% NaCl and resuspended in 1.5 mL lysis buffer (25 mM Tris-HCl, pH 8.0, 100 mM NaCl, 2 mM EDTA) followed by incubation with 0.2 g/L lysozyme on ice for 45 min and sonication (3 min, 35% duty cycle, 3 output control). After addition of 10 mM MgCl2 and treatment with 125 U Benzonase Nuclease (Sigma-Aldrich) for 10 min, the lysate was centrifuged to separate the soluble supernatant fraction from the pellet. The insoluble pellet fraction was resuspended in 1.5 mL lysis buffer. Both fractions were run on an SDS-PAGE using 12% ClearPageTM gels and ClearPAGETM SDS-R Run buffer (C.B.S. Scientific) followed by staining with Coomassie Brilliant blue R-250 (Merck). Western blot analysis was performed as described elsewhere^33^.

### mCherry production analysis

mCherry activity was determined with an Infinite M200 Pro multifunctional microplate reader (Tecan) by measuring the fluorescence of 100 μL untreated culture with excitation and emission wavelengths of 584 nm (9 nm bandwidth) and 620 nm (20 nm bandwidth), respectively, and normalisation against absorbance reading at OD_600_. Three bacterial colonies from each of the dual UTR constructs were transferred into a 96-well plate (Corning, black, flat bottom, clear) with each well containing 100 μL LB with kanamycin. *E. coli* clones were grown until they reached mid log-phase and were induced either with L-arabinose to a final concentration of 0.2% or with m-toluic acid at concentrations specified. The induced cultures were grown for 5 and/or 18 hours (O/N) as specified in the main text.

### Transcript analysis

To relatively quantify *bla* and *mcherry* transcripts, *E. coli* RV308 clones were cultivated as described above. For RNA decay studies, *E. coli* clones with dual UTR constructs were grown until the cultures reached the mid-log phase and were harvested 5 hours after induction with 2 mM *m*-toluic acid. Samples for transcript analysis were taken 2, 4, 6, 8, and 10 min after filtration following the wash-out protocol^32^. Cell cultures were treated with RNA protect (Qiagen) to stabilise RNA. RNA isolation, DNase treatment, cDNA synthesis, and qRT-PCR were performed according to protocols described elsewhere^18^. The following primer pairs were used in qPCR: for *bla* (5′-ACGTTTT CCAATGATGAGCACTT-3’and 5′-TGCCCGGCGTCAACAC-3’); for mCherry (5′-CGTTCGCTTGGGACAT CCT-3’and 5′-GATGTCAGCCGGGTGTTTAAC-3’, and for 16S rRNA (5′-ATTGACGTTACCCGCAGAAGA A-3’and 5′-GCTTGCACCCTCCGTATTACC-3’).

### RBS calculator details

Translation initiation rates were determined using the reverse engineering function of the RBS calculator^24^. The sequence input for the RBS calculator consisted of the 5′ UTR DNA sequence (up to 50 nt) and the first 50 nt of the *bla* or *mcherry* coding sequence. 5′ UTRs with optimal translational features were generated applying the forward engineering function of the RBS calculator using the following template sequence 5′-GAGCTCCATTATTATTGT ATATGTnnnnnnnnnnnnnnnnnnnnnnnnT-3’.

## Supporting information

Plasmid maps

## Acknowledgements

Authors would like to thank Laila Berg, for giving valuable technical advice for the library construction and screening; Svein Valla (deceased), for his involvement in an earlier version of the manuscript.

This work was supported by the Faculty of Natural Sciences at the Norwegian University of Science and Technology, Trondheim, Norway (personal PhD stipend that SB received) and the Norwegian Research Council grant numbers 192432 and 192123.

## Author contributions statement

RL conceived the study. SBL and RL designed the experiments. SBL, IO and JAL carried out the experiments. All authors analysed the results. SBL and RL wrote the manuscript with the help from co-authors. All authors reviewed the manuscript.

## Additional information

The dual UTR concept has been patented (Application granted on July 24^th^, 2019).

