## Supporting Online Material for "Dual UTR-A novel 5′ untranslated region design for synthetic biology applications"

Le et al. 2019

**Table S1: Nucleotide sequence composition of the Tr- and Tn-UTRs identified in pAO-Tr- and pAO-Tn-based libraries and the ampicillin resistance phenotype of the *E. coli* clones harbouring the constructs.**

The values depicted correspond to maximal ampicillin concentrations [g/L] at which growth was observed. LV-2 is a 5′-UTR variant identified by Berg et al. 2009. The PciI recognition sequence is underlined, the minimal Shine-Dalgarno sequences are double-underlined, and the ATG start codon is typed in boldface. Numbers in brackets indicate the next ampicillin concentrations tested above the tolerated maximum.

| Plasmid | Sequence 5′ -> 3′ | *m*-toluic acid concentrations | |
|---|---|---|---|
| | | 0 mM | 0.1 mM |
| **pAO-Tr** | | | |
| wt | AACATGT-ACAATAATAATGGAGTCATGAACAT**ATG** | 0.025 (0.050) | 0.25 (0.40) |
| LV-2 | .......-..C......CA.....T........... | 0.025 (0.050) | 1.0 (1.2) |
| r11 | .......-..C.......C................. | 0.015 (0.025) | 0.60 (0.8) |
| r28 | .......-.-T............AA........... | 0.015 (0.025) | 0.80 (1.0) |
| r31 | .......T..C..G...................... | 0.025 (0.050) | 1.0 (1.2) |
| r36 | .......-....GT....C.....A........... | 0.025 (0.050) | 1.0 (1.2) |
| r50 | .......T..........C.........T....... | 0.025 (0.050) | 1.0 (1.2) |
| **pAO-Tn** | | | |
| wt | AACATGTACAATAATAATGGAGTCATGAACAT**ATG** | 0.010 (0.025) | 0.10 (0.25) |
| n2 | ........GTT.....-.........T........ | 0.25 (0.50) | 2.0 (2.5) |
| n3 | .........T.A.C........AA........... | 0.25 (0.50) | 2.5 (3.0) |
| n13 | .......G..................C........ | 0.25 (0.50) | 2.0 (2.5) |
| n15 | ...........C....G.........T........ | 0.25 (0.50) | 2.0 (2.5) |
| n16 | ............C.............A........ | 0.25 (0.50) | 1.5 (2.0) |
| n17 | .......G..................T........ | 0.25 (0.50) | 2.0 (2.5) |
| n18 | ........A..A.G............T........ | 0.25 (0.50) | 2.0 (2.5) |
| n24 | ..........T.....TA........C........ | 0.25 (0.50) | 2.0 (2.5) |
| n15 | ..........C.....G......T........... | 0.25 (0.50) | 1.5 (2.0) |
| n17 | ........A.C...............T........ | 0.25 (0.50) | 2.0 (2.5) |
| n23 | .............G............T........ | 0.25 (0.50) | 2.0 (2.5) |
| n25 | ............G.............A........ | 0.25 (0.50) | 1.0 (1.5) |
| n35 | ........A.......TA........C........ | 0.25 (0.50) | 1.5 (2.0) |
| n39 | ........-.................T........ | 0.25 (0.50) | 2.0 (2.5) |
| n41 | ........A..C..C..C........T........ | 0.25 (0.50) | 2.5 (3.0) |
| n42 | ........A.....CT..........A........ | 0.25 (0.50) | 2.0 (2.5) |
| n44 | .......G.........A........C........ | 0.25 (0.50) | 2.5 (3.0) |
| n47 | ........AT.A.C...A.....T........... | 0.25 (0.50) | 2.5 (3.0) |
| n48 | .........T...T........AG..T........ | 0.25 (0.50) | 2.0 (2.5) |
| n52 | ........GT...GA...........T........ | 0.25 (0.50) | 2.0 (2.5) |
| n58 | ..........T...C..A........AT....... | 0.25 (0.50) | 2.5 (3.0) |
| n59 | .........T..T.G.TA........T........ | 0.25 (0.50) | 2.5 (3.0) |

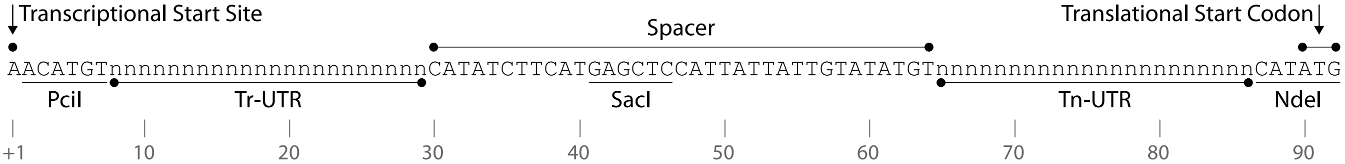


**Figure S1: Nucleotide sequence composition of dual UTR.**

The transcriptional start site (A) is indicted with +1. The restriction enzyme recognition sequences are underlined, and Tr- and Tn-UTR and spacer regions are marked.

**Table S2: The sequence composition of the RBS calculator designed 5′-UTR DNA sequences.**

Template is the sequence used as a template in the design of 5′-UTR DNA sequences, the putative Shine-Dalgarno sequences are double-underlined and the ATG start codon is typed in boldface. dTn1-6 represent the sequence composition of designed six Tn-UTRs with maximal TIRs with *bla* (dTn 1-3) or *mcherry* (dTn 4-6) coding sequences.

| Name | Sequence 5′→ 3′ ---------------------n24---------------- |
|---|---|
| Template | GAGCTCCATTATTATTGTATATGTnnnnnnnnnnnnnnnnnnnnnnnnT**ATG** |
| dTn1 | ........................GCATCAATTACTAAGGAGGTATACT... |
| dTn2 | ........................GCATCACCCTTTAAGGAGGTTTACT... |
| dTn3 | ........................ACCGTACCCGTTAAGGAGGTTTTCT... |
| dTn4 | ........................AACAAGGCAGAATAAGGAGGTTCAT... |
| dTn5 | ........................GGATATACCCAGTAAGGAGGTACAT... |
| dTn6 | ........................ATATAAGGATTAGAGGAGGTAATAT... |

**Table S3: Calculated translation initiation rates of Tn-dual UTRs in combination with *bla* and *mCherry* coding sequence.**

dTn1-6 represent sequences of six Tn-UTRs with maximal TIRs for *bla* (dTn1-3) or *mcherry* (dTn4-6). The TIR for the Tn-UTRs wt, n24, n44, n47 and n58 were calculated using the reverse engineering function of the RBS-calculator.

| | TIR | |
|---|---|---|
| **Tn-UTR** | ***bla*** | ***mCherry*** |
| wt | 598.4 | 2,308.6 |
| n24 | 5,332.0 | 5,678.7 |
| n44 | 7,994.5 | 25,075.0 |
| n47 | 5,834.1 | 12,766.2 |
| n58 | 3,461.4 | 4,743.2 |
| dTn1 | 349,161.5 | - |
| dTn2 | 418,029.8 | - |
| dTn3 | 478,456.9 | - |
| dTn4 | - | 856,820.0 |
| dTn5 | - | 819,114.1 |
| dTn6 | - | 655,630.7 |
